## Supplementary figures and images for "Viral-mediated Oct4 overexpression and inhibition of Notch signaling synergistically induce neurogenic competence in mammalian Müller glia"

### Figure S1

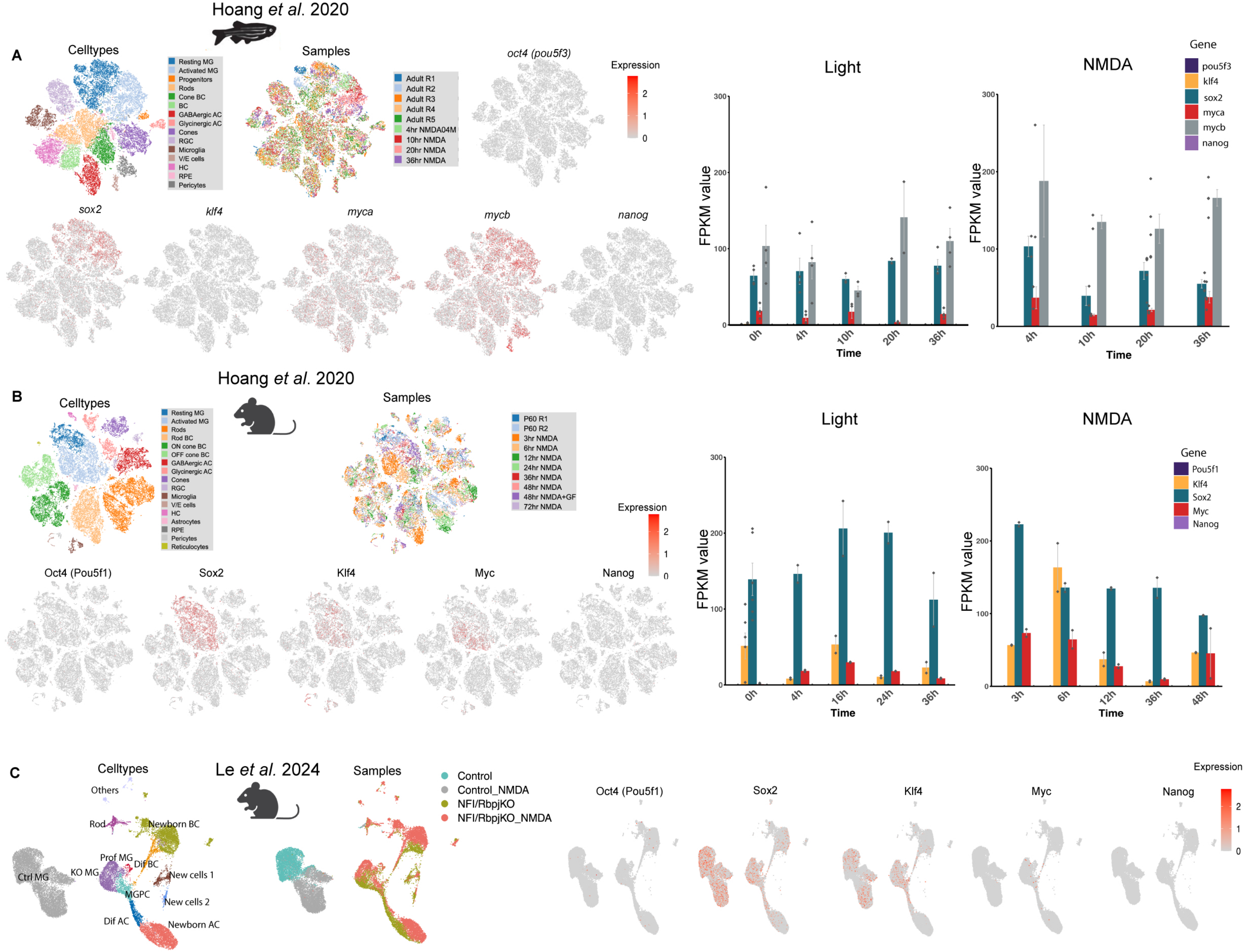


Le, et al. Fig. S1

### Figure S2

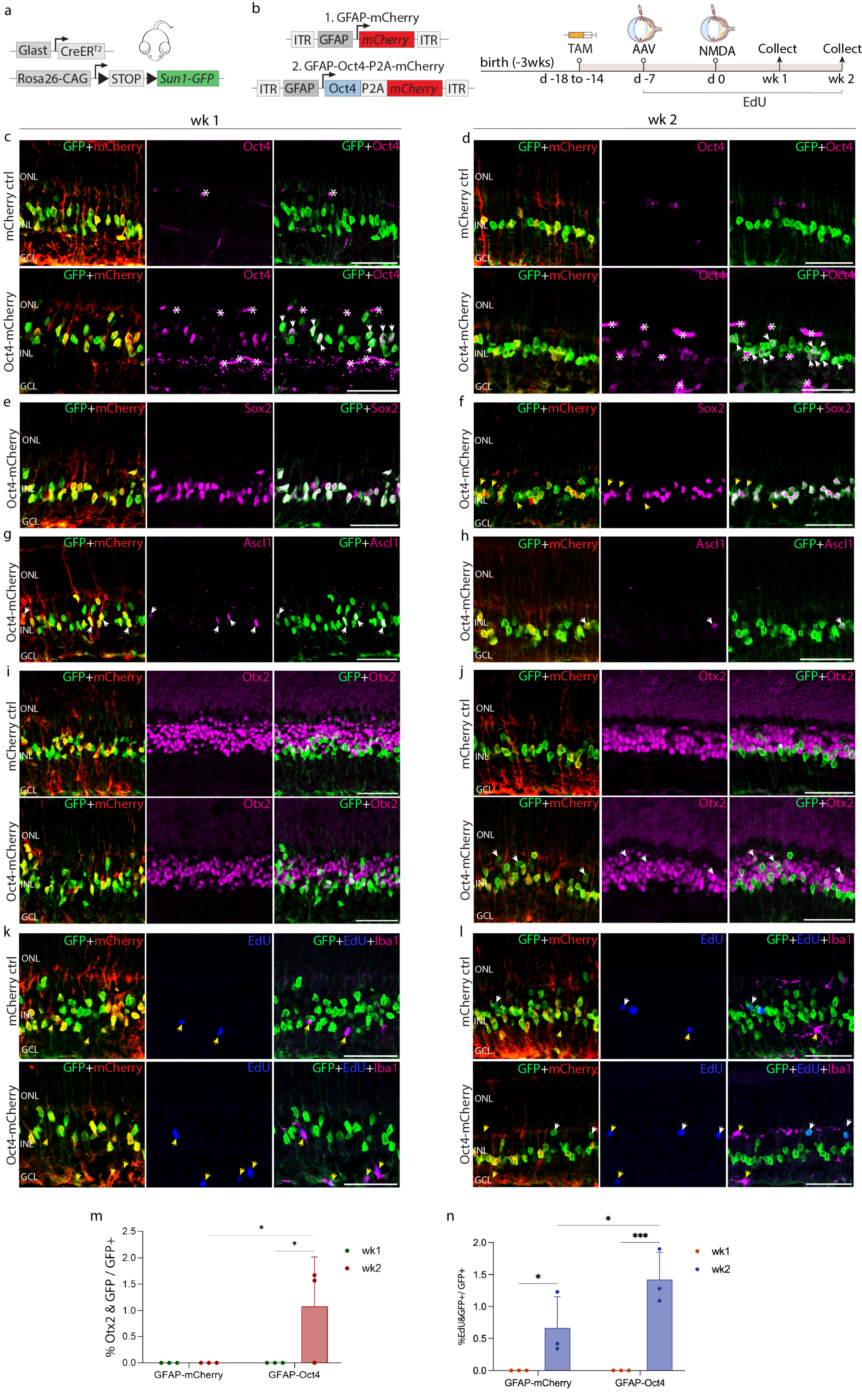


Le, et al. Fig. S2

### Figure S3

*GlastCreER;Sun1-GFP* (5 wks post NMDA)

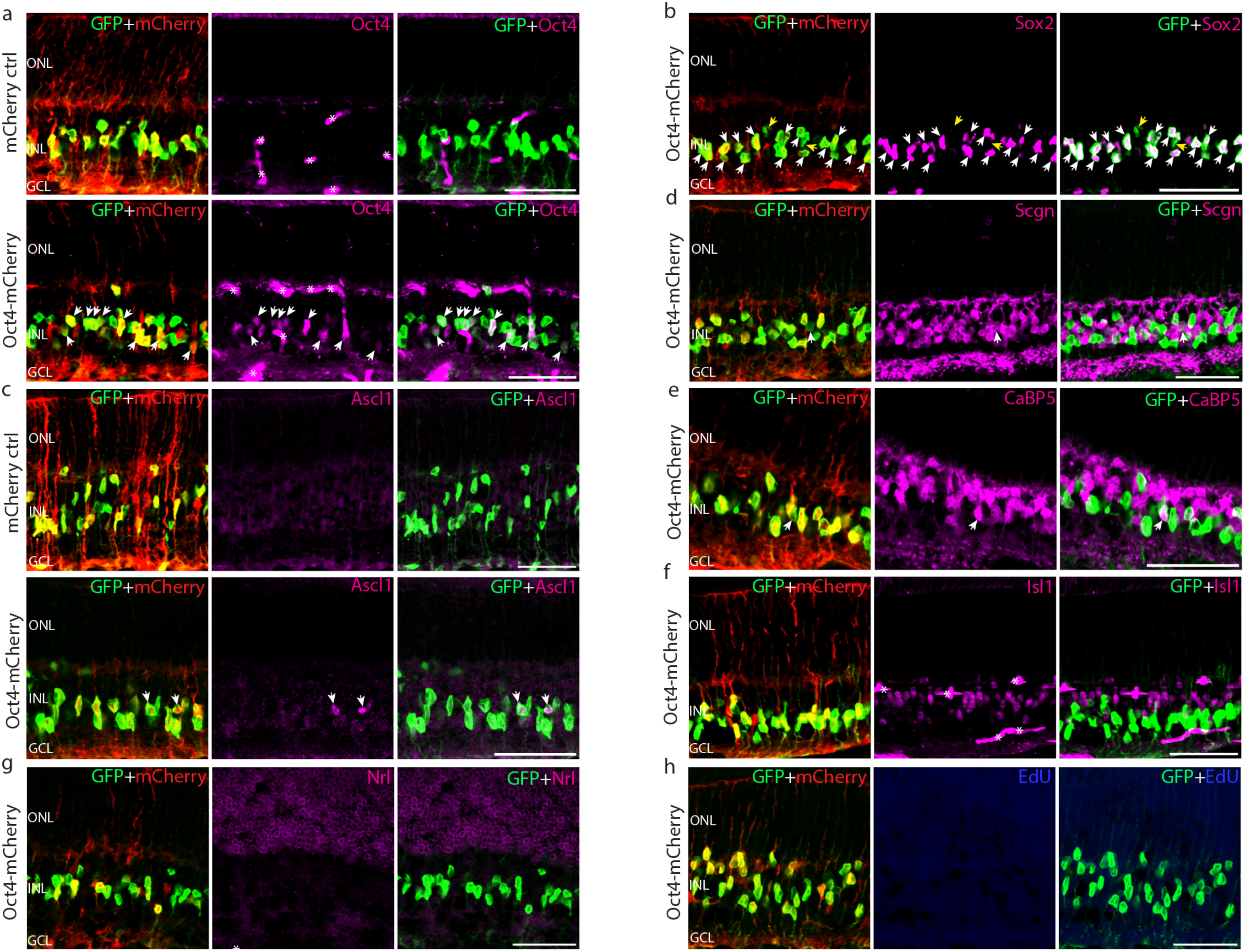


Le, et al. Fig. S3

### Figure S4

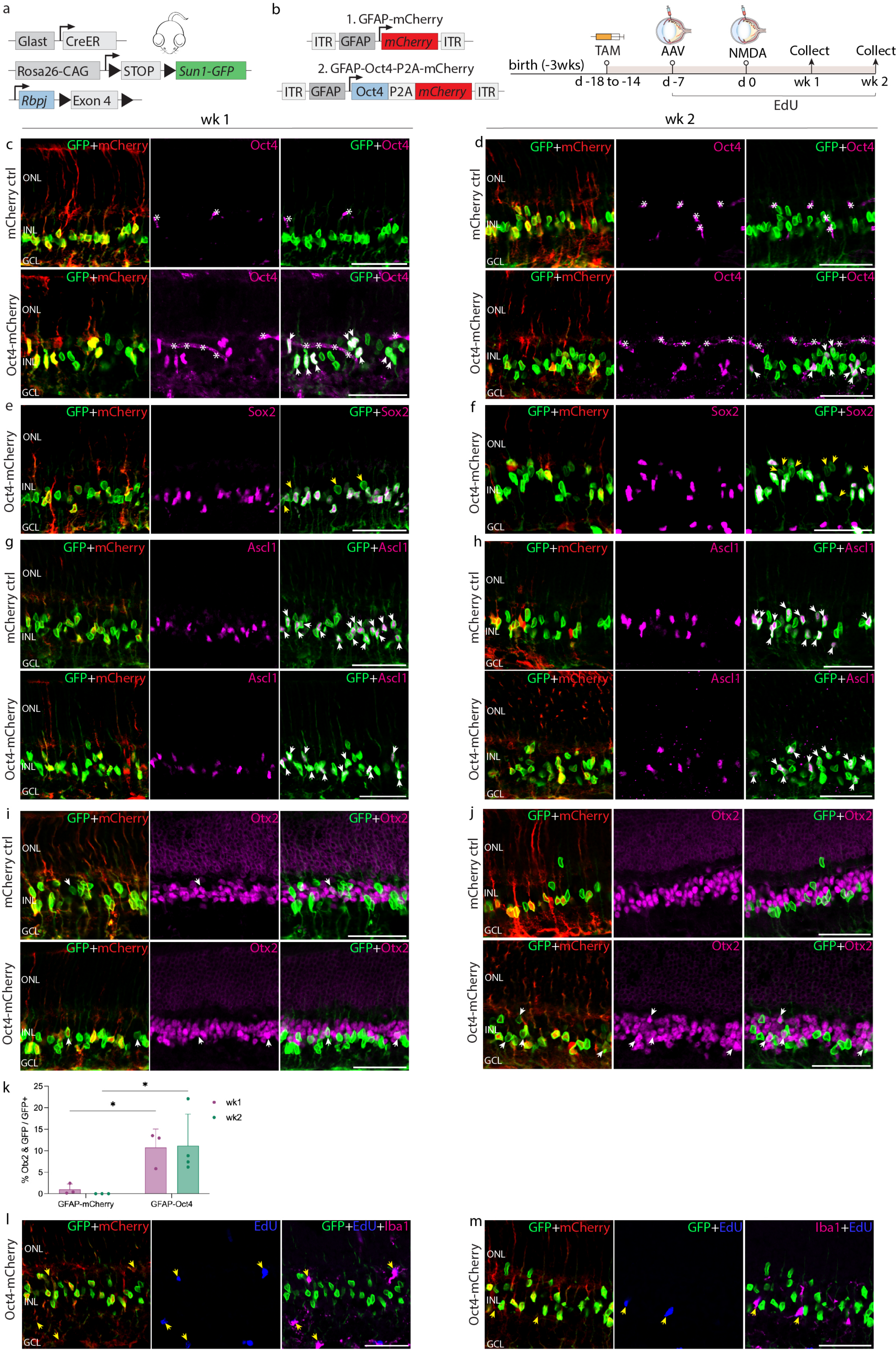


Le et al. Fig. S4

### Figure S5

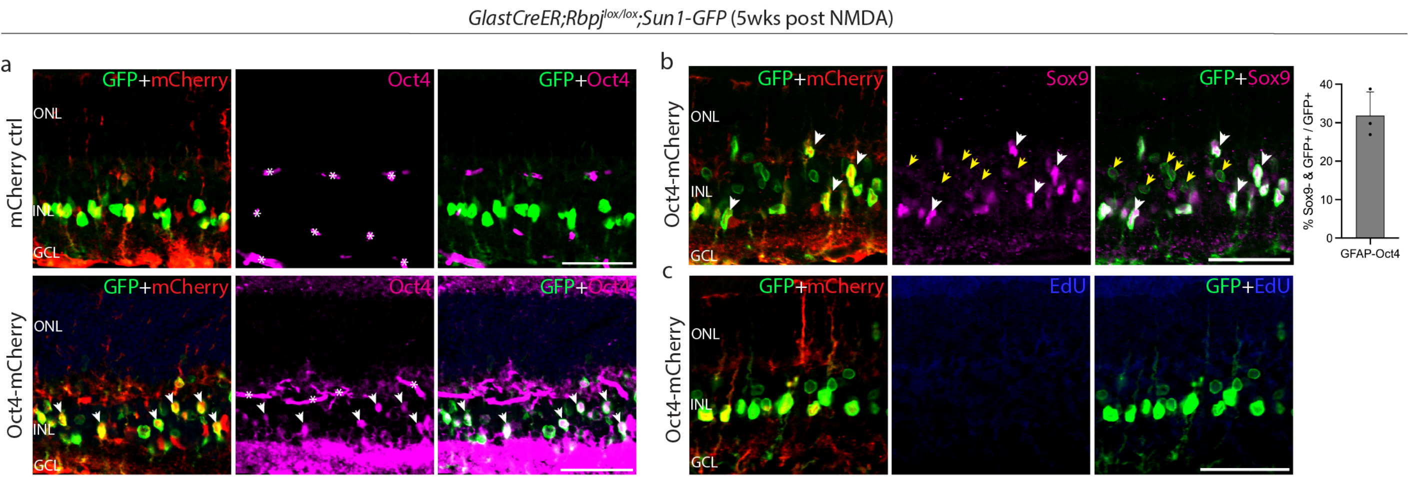


Le, et al. Fig. S5

### Figure S6

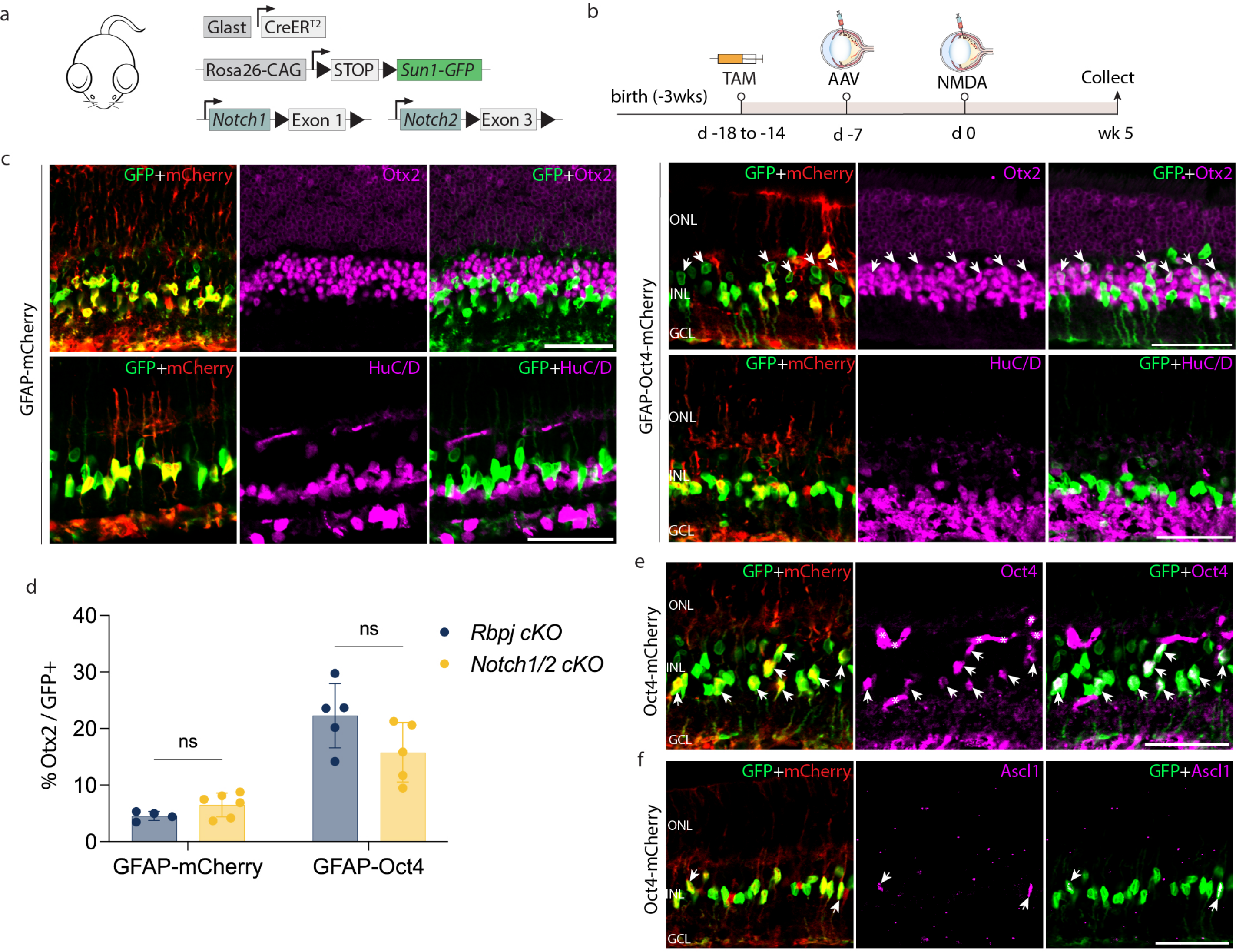


Le, et al. Fig. S6

### Figure S7

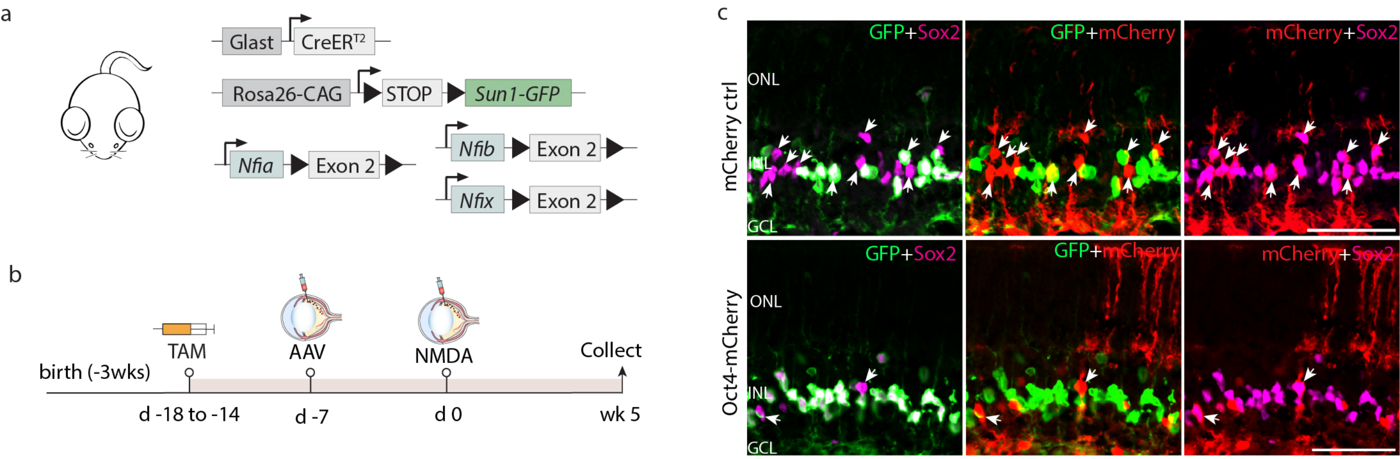


Le, et al. Fig. S7

### Figure S8

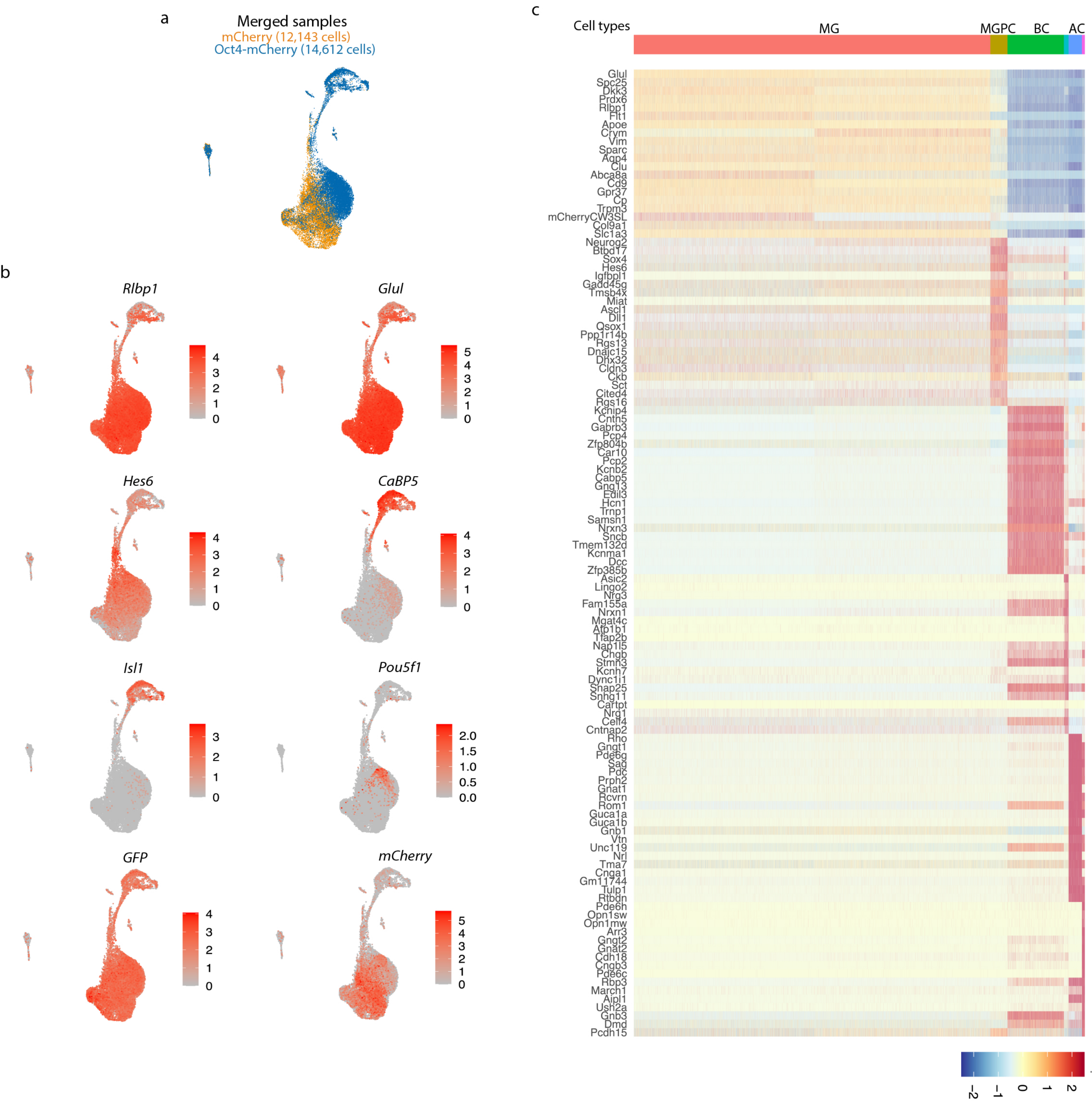


Le, et al. Fig. S8

### Figure S9

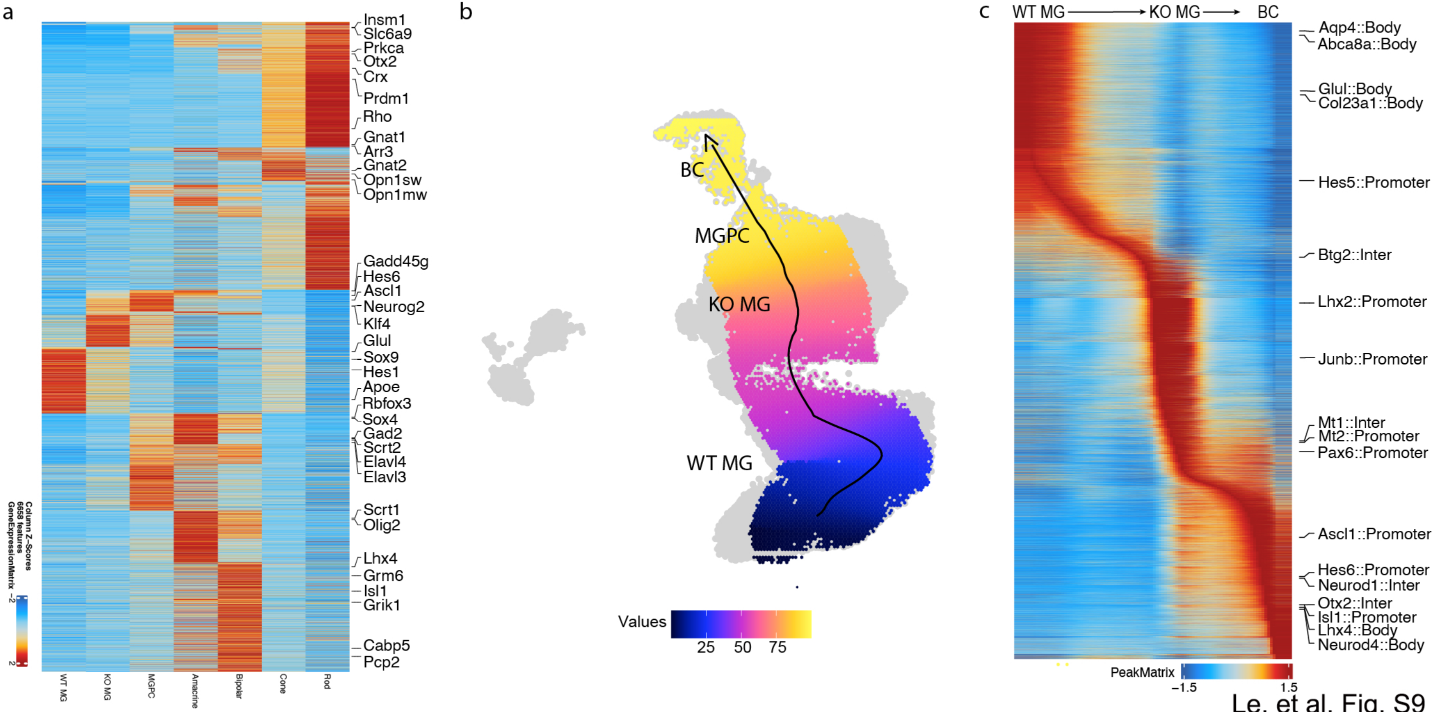


Le, et al. Fig. S9
